# Three-dimensional localization of fluorescent proteins in living *Escherichia coli*

**DOI:** 10.1101/2023.10.20.563292

**Authors:** Praneeth Karempudi, Konrad Gras, Elias Amselem, Spartak Zikrin, Johan Elf

## Abstract

3D localization of fluorescent proteins (FPs) in living bacteria has been challenging due to the low signal-to-background ratio of the FPs and the relatively uncertain positioning of the cells in the optical reference system. Using mother-machine microfluidic devices together with deep learning, we present an approach that enables accurate 3D localization of FPs in *Escherichia coli* over long periods. We describe a method to simulate ground truth training data for the deep learning network based on background models generated from experimental data. We test the method by studying how chromosomal loci are relocated in 3D over the *E. coli* cell cycle. Since the cells are radially symmetric, we expect the same width and height distribution of fluorophores if the 3D positions are correctly determined. We observe this pattern experimentally for all the labelled loci on the chromosome. Interestingly, some loci are located exclusively in the periphery of the nucleoid, while others are more confined to the core of the nucleoid. This method enables studying any chromosomal loci inside living *E. coli* cells in high-throughput.

## Introduction

Single-molecule localization microscopy (SMLM) is widely used in biology to resolve cellular structures below the diffraction limit, characterize molecular movement, and study spatial distributions of proteins and macro-molecules [1]. Structures of interest are typically labeled with fluorophores and imaged using a wide-field fluorescence microscope equipped with appropriate excitation and emission filters. In super-resolution methods like PALM[2] and (d) STORM[3, 4], cells are usually fixed, and fluorescence imaging is done in multiple cycles by exciting and localizing only a few emitters in each cycle. Emitter locations are pooled from all cycles to render a final high-resolution image. The density of emitters, exposure times, and frame rates are optimized together with the localization algorithms to resolve structures with nanometer precision [5].

SMLM has also been used to study molecular diffusion properties and their state-transitions in living cells [6–8]. Molecules of interest are labeled with bright dyes and imaged in time-lapse to construct molecular motion trajectories for further analysis. These studies typically involve a small number of cells where molecules are tracked over a few tens of frames, with short exposure times as well as shorter intervals between consecutive camera frames matching the timescale of molecular diffusion. However, imaging FPs in living cells over longer experimental time frames requires trade-offs between localization accuracy, time resolution, exposure times, and bleaching. In bacterial cell biology, imaging has advanced to a stage where we can track intracellular structures over many generations to address questions related to, for example, coordination of growth, division, and replication. These experiments are typically performed in microfluidic devices, specifically mother-machine devices [9, 10]. Mother-machines offer the ability to image thousands of individual bacteria for many generations, confined to a single imaging plane. As cells grow and divide, extrinsically supplied cellular labels get diluted. For this reason, imaging on the cell cycle time scale typically relies on intrinsically expressed fluorescent proteins (FPs) that get renewed as the cells grow. The main problem with FPs is that they are not very bright and unbound emitters can contribute to the background fluorescence signal, which makes it challenging to accurately localize single emitters.

Precise localization of emitters from imaging data is critical in SMLM. Methods for localization of molecules in 2D (Gaussian PSF) using maximum-likelihood estimation formulations [11–13], and their limits have been well-studied [14, 15]. SMLM in 3D can be achieved by encoding the z-position of a fluorescence emitter into the point spread function (PSF) to break its symmetry along the z-axis [16]. Popular PSF-engineering methods include the astigmatic PSF [17], double-helix PSF [18], saddle-point PSF [19], and the tetrapod PSF [20]. As our applications are usually limited by emitters with low photon counts and multiple emitters inside a single cell, spreading the PSF over a large area or losing photons to transmission-reducing optical components would result in a very low signal-to-background ratio. We are also interested in a limited z range defined by the width of a bacterial cell. In this study, we therefore use the astigmatic PSF to localize emitters.

Theoretical models of measured PSFs might not capture all optical and system specific aberrations, so we instead use experimentally measured PSFs (SI A.3) that are parameterized using spline models. Recent methods show that experimental PSFs can be interpolated with splines [21–23], which can then be used for fitting emitter location using optimization algorithms. GPU implementation of fitting algorithms [24] reduces computation time significantly, enabling analysis of larger datasets, fitting as many as *>* 10^5^ emitters/second.

Deep learning-based algorithms DeepSTORM3D [25], DeepLoco [26], and DECODE [27] successfully apply convolutional-neural-networks (CNNs) to achieve improved localization precision in both single-emitter and multi-emitter regimes. CNNs performs well under a wide variety of conditions, with the requirement that the simulated data used to train the neural network represents realistic data sets. In a typical field-of-view (FOV), background structures that are not labeled with fluorophores also contribute signal to the image and need to be included in the training data. The background photons can be sampled from a uniform distribution over a range of values [27] or modeled as diffuse structures using multi-scale Perlin noise models [28]. The use of CNNs for structured background removal has been shown to improve localization precision [28]. For FPs expressed in *E. coli* cells growing in mother-machine devices, the size of the cell contributing to background photons is the same as area over which the PSF (Fig 1b) is spread on the image. Hence, accurate modeling of the background is required to achieve good localization of point emitters.

**Fig. 1.**
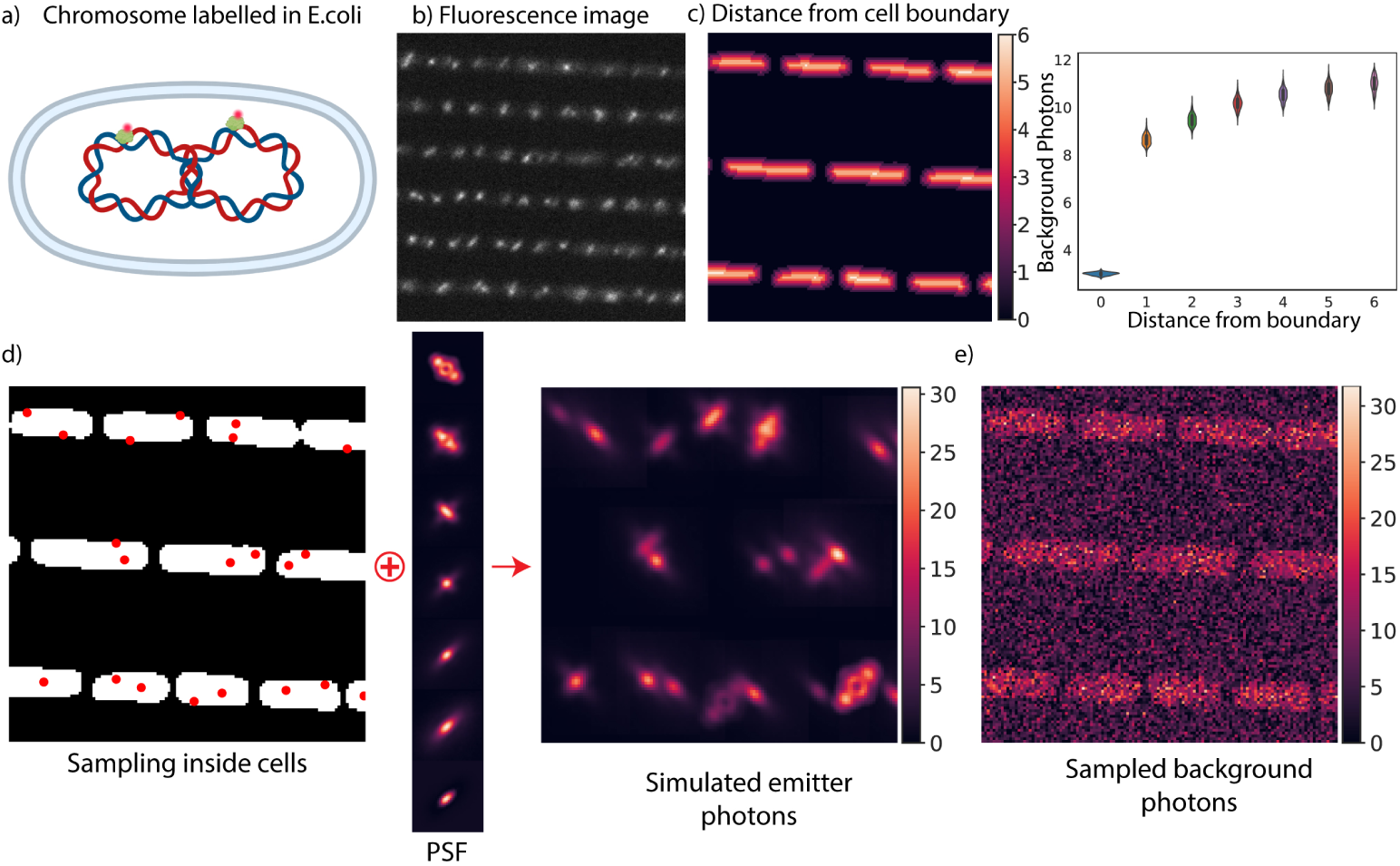
a) Illustration of an *E. coli* chromosome labeled with fluorescent proteins b) YFP fluorescence image of cells growing in the mother-machine device. In the image, we can see the emitters and the structured cell background c) Left: Distance from the cell boundary in pixels calculated using the Euclidean distance transform on the cell segmentation of a phase-contrast image. Right: Background photons distribution over time as a function of distance from the cell boundary. Each data point at a particular distance is the mean background photons from all the cells in a single image. The curve shows a consistent background profile for the duration of the experiment containing 300 frames. d) Left: Sampled emitter locations inside cells (red dots) overlaid on a cell segmentation mask. Middle: Spline-interpolated PSF sampled at different z (-700 nm to 700 nm). Right: Emitters with photon counts between 750 and 3000 sampled from the PSF spline model. e) An image showing background photons sampled from the background model.

3D localization of fluorescent proteins in live *Escherichia coli* cells hasn’t been explored well due to the variable background noise and low signal-to-background ratio. In this work, we address the challenge of localizing FPs in *E. coli* in 3D using deep learning and data-driven simulations. We use a mother-machine-based fluidic device with open-ended growth channels to trap and image growing *E. coli* for extended periods. To train the neural network models, we need simulated images representing accurate background structures and emitters similar to experimental data, with known emitter properties. The simulations require a model of the cell background, emitter PSF and camera noise. We simulate emitters using a cubic spline model of the PSF and approximate the background structures using experimental data. Background photons varying across the FOV are estimated by fitting appropriate probability distributions to experimental data. These distributions are used for sampling during training of the neural network model. We also perform camera calibration of our sCMOS camera (Methods 1) to obtain pixel-dependent camera noise that we use for training data simulations. In summary, we develop a high-precision 3D localization algorithm using a field-dependent DeepLoc (FD-DeepLoc) [29] approach to encode sub-locations of the camera chip together with cell segmentation and estimation of background photon patterns. Finally, we apply this localization algorithm to localize positions on the chromosomal DNA and the replisome complex throughout the *E. coli* cell cycle.

## Results

### 3D fluorescent emitter imaging in live *E. coli*

We imaged live *E. coli* cells over several division cycles in a microfluidic mothermachine-type device [9], where a constant supply of growth medium was maintained to support exponential growth and long-term imaging. Phase-contrast images were acquired every minute to monitor the growth of the cells. We engineered *E. coli* strains with labels on the replication complex as well as on different chromosomal loci (Fig 1a). Three loci on the chromosome were chosen to be labeled, one near the origin (*oriC*), one near the terminus (*ter*), and one midway between the origin and terminus (midway). Locus labeling was achieved by fusing a yellow fluorescent protein (YFP) with the DNA-binding protein MalI [30]. An array of twelve MalI-binding operator sequences was introduced in different locations on the chromosome. The replisome was labeled using a translational fusion between DnaN and a YFP [31]. The fluorescent labels were imaged every 2 min with a custom optical setup [30] equipped with an astigmatic lens in the optical path (Methods 1). Example images of fluorescent emitters are shown in Fig 1b. Phase-contrast images were segmented with the Omnipose cell segmentation algorithm described in [32, 33]. The segmentation masks were also used in the training-data simulation for the localization neural network.

### Background estimation and generation of simulated images

Accurate estimation of background has been previously shown to improve localization precision [28]. The structured background observed in live bacteria (Fig 1b) requires a different approximation compared to [27, 28] due to similar spatial scales of background structures and the point spread function of the emitters. Fluorescence images were thresholded to remove most of the emitters’ signal from the image (SI Fig A4). Using the Euclidean distance transform (EDT) of the cell segmentation mask, we aggregated the photons produced by the cell background as well as the fluidic chip (Fig 1c, left). The distributions of photons at different distances from the boundary were modeled using a gamma distribution. The means of the gamma distribution over 300 images follow a pattern shown in Fig 1c(right).

Neural network-based localization algorithms use simulated images of fluorescent emitters as training data. Images of emitters need to be generated using an accurate model of the PSF. We used cubic splines to parameterize the shape of the PSF [23, 24] and to sample images of emitters in 3D at specific locations and photon counts (Methods 1). Background photon distributions were approximated from experimental data for the chromosomal dots (SI A.5) and the replisome dots (SI A.6). To generate simulated images, we sampled emitter location pixels on cell segmentation masks with a fixed average emitter density per cell (Fig 1d). Sub-pixel offsets of the emitters in x and y from the center of their pixel were sampled uniformly within a one-pixel range. Z coordinates of emitters were sampled from a uniform distribution that covered the experimental imaging range. The cubic spline PSF model with the emitter locations incorporated was used to simulate fluorescence emitters of chromosomal loci (Fig 1d). Cell background photons sampled from one such background model are shown in Fig 1e to illustrate the diffuse features, closely resembling the features found in experimental data. Emitter sampling and simulation image generation are described in detail in the SI section A.4.

Images of sampled emitters were overlaid on backgrounds sampled using EDT-based distributions to obtain training data in photon units. These images were passed through the sCMOS camera noise model (SI section A.1,A.2) to generate final images in analog-to-digital units (ADU) (SI Fig A3f). The parameters used for simulations, such as emitter photon count range, background distribution, and emitter density in cells were adjusted to represent the experimental data to which the model was applied.

### Emitter localization using a deep neural network

We used the simulated images containing cells with emitters and realistic background noise as training data for the 3D localization algorithm. Sub-regions (128 x 128 pixels) (Fig 1d-e) of the full camera FOV (1302 x 1041 pixels) together with the corresponding sub-region of the normalized XY positions on the camera were used to train a FD-DeepLoc model [29] (Fig 2a) until convergence (SI A.8). This neural network takes an input sub-region image together with two images encoding the XY positions of the pixels of this sub-region on the full camera FOV and produces ten output images, which encode the locations, photon counts, and their uncertainties. The architecture of the network, encoding of the positions as network outputs, training, and loss functions, are described in the Methods section 1. Simulated images of the same size as the subregions with known emitter locations were randomly generated during the training process. Evaluation of the model was done on 30 full-camera FOV images containing ~ 20,000 emitters sampled at the beginning of training. Each model was evaluated with performance metrics from the SMLM challenge 2016 [34] (SI A.8) on these 30 images. To evaluate model performance, predicted emitters were matched to ground truth emitters using criteria such as probability predicted by the network and Euclidean distance in 3D. The training for the two scenarios, chromosomal loci labels or replisome labels, only differed in terms of background noise model (SI A.5, A.6). Models that achieved high precision (~1.0) and recall (~0.8-0.9) and low RMSE in x, y, and z were trained for both chromosomal loci and replisome labels. Distributions of x, y, and z, photon counts, and uncertainty measures predicted by the neural network are described in SI section A.8.

**Fig. 2.**
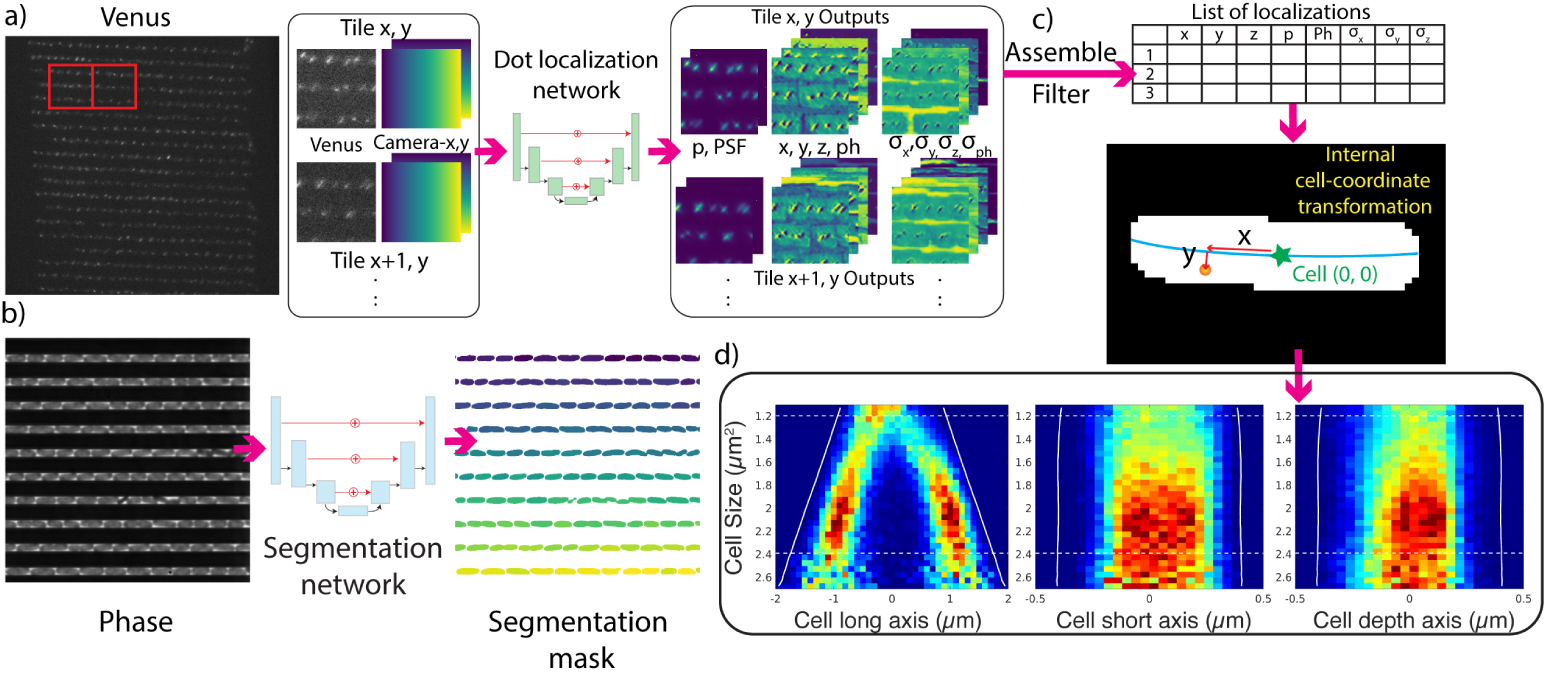
a) Localization of emitters using a deep neural network. The fluorescence image is chunked into tiles together with the camera location field and fed to a neural network. The network outputs 10 images for each tile in the large image (one for the probability of finding an emitter in a pixel (p), one for the cell-background-free PSF (PSF), four images for sub-pixel-x, y, and z (x, y, z), and photon counts (ph), and four for their corresponding precision predictions (*σ_x_, σ_y_, σ_z_, σ_ph_*). b) Phase-contrast image segmented using the Omnipose neural network to produce segmentation labels for each cell. c) Top: Localization network outputs from each tile of the fluorescence channel image are assembled and filtered to generate the final coordinates of the emitters with respect to the origin of the image. Bottom: Emitters are assigned to cells using a cell segmentation mask and their coordinates are converted to the cells’ internal coordinates with respect to their origin (green star) and backbone (blue curve). d) Localizations from time-lapse imaging are pooled together, binned by cell size, and plotted as the distribution of emitters over the cell cycle. White lines indicate the average position of the cell poles, for the cell long axis figures, or the cell widths, for the cell short and depth axes figures.

Phase-contrast images acquired in conjunction with the fluorescence images were segmented using an Omnipose cell segmentation network (Fig 2b). The trained neural network models were applied to experimental images of different loci to obtain a list of emitter coordinates. Transformation of emitter coordinates onto the corresponding phase-contrast image was required as these images were captured on two different cameras (SI section A.9). After this transformation, emitter positions in each image were overlaid on the corresponding cell segmentation mask, and XY coordinates of emitters falling inside the cells were converted to internal long- and short-axis coordinates, with the origin of a cell being the center of its long-axis backbone and all cells being oriented from old pole to new pole (Fig 2c).

The internal coordinates in x and y can be obtained using the method described above. However, the internal z coordinate cannot be assigned due to a lack of a reference origin on the z-axis. The z predictions from the localization network are relative to the zero of the PSF (SI A.3). The mean values of predicted z coordinates across the FOV were fit to a 2D plane and localizations were offset in z using the values of z on this plane. This correction accounts for z variations induced by the fluidic chip placement on the microscope and centers the depth (z)-axis localizations around zero defined for each point by this plane. The boundaries of the cell in the depth axis are however reflected on the y-axis due to the radial symmetry of the *E. coli* cell around its long axis. Emitter locations from cells imaged over several cell division cycles were pooled to obtain position distributions in x, y, and z as a function of cell size (Fig 2 d).

### Consistent 3D localization of fluorescent emitters across the cell cycle

We investigated the positions of the bacterial replisome and several chromosomal loci as a function of cell size. 3D position distributions based on the output from the neural network were visualized as heat maps over the cells’ long, short, and depth axes in (Fig. 3).

**Fig. 3.**
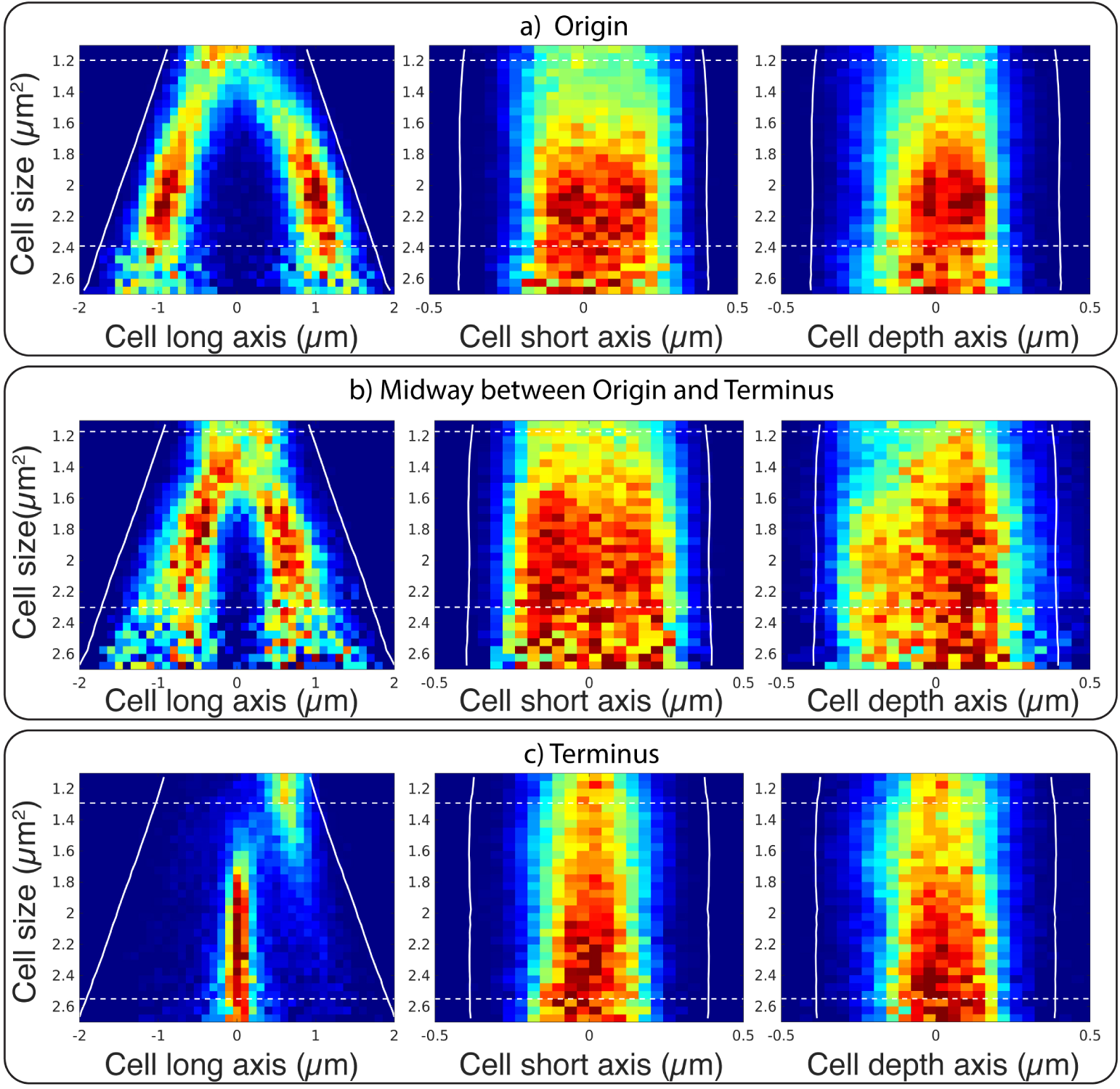
Localization of different loci on the *E. coli* chromosome as a function of cell size. Labelled loci: a) Origin b) Midway between origin and terminus c) Terminus. The x-axis for each figure represents the cell long axis or x-coordinate (left), cell short axis or y-coordinate (center), and cell depth axis or z-coordinate (right). The y-axis represents cell size in *µm*^2^. The positions of each loci are pooled from 20 mother-machine traps with growing cells imaged every 2 minutes for 600 minutes.

The long (x-)axis and short (y-)axis location distributions for all loci and the replisome are similar to previously reported location patterns ([30, 31, 35, 36]). The positions on the depth (z-)axis dimension are less straightforward to corroborate since they have not been observed before. However, based on the radial symmetry of *E. coli* cells, we expect that the position distributions in y and z should be identical. In line with this assumption, the measurements do confirm that the distributions of y and z coordinates show a large degree of similarity. When the distributions are plotted in the y-z plane (Fig 4), all loci exhibit distinct radially symmetric location patterns, which further supports that we can accurately localize emitters on the z-axis.

**Fig. 4.**
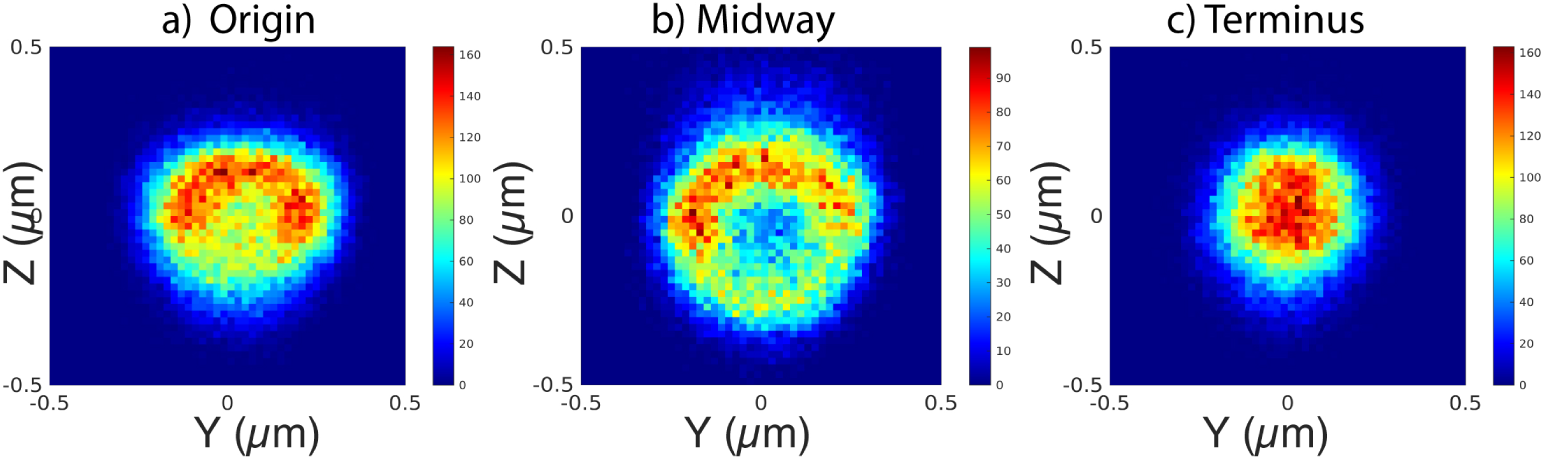
YZ distributions of the location of different loci a) Origin b) Midway between origin and terminus c) Terminus. Bin size is 20 nm x 20 nm. Colorbars show the number of localizations in each bin.

The *ter* label has a relatively confined localization in the cell center in all dimensions. In contrast, the midway locus at 1.1 Mbp from *oriC* has a bimodal short-axis localization distribution throughout the division cycle, which we also observed even more clearly in the z distribution. This radial pattern implies that the locus is located close to the edge of the nucleoid, possibly due to interactions with the cell membrane [37] or proximity to highly expressed genes [38]. The ring-like distribution gives us an opportunity to estimate the upper bound of the combined errors from localizing the FPs and pooling data from different cells. If we assume that the real localizations lie on a perfect circle and are the same in all cells, the broadening of the circle realized as a ring in Fig 4b, implies that the standard deviation of experimental error is at most 50 nm. The *oriC* label exhibits a relatively broad radial distribution that differs in its shape compared to both distributions for the *ter* and midway labels.

The localization of the replisome complex has previously been described as being confined [39], a pattern that we also capture in our results (Fig 5). By visualizing how the 2D positions of *oriC* and the replisome change throughout the cell cycle, we recently showed that replication initiation does not occur at the cell membrane [30]. The radial position distributions presented here further support this conclusion.

**Fig. 5.**
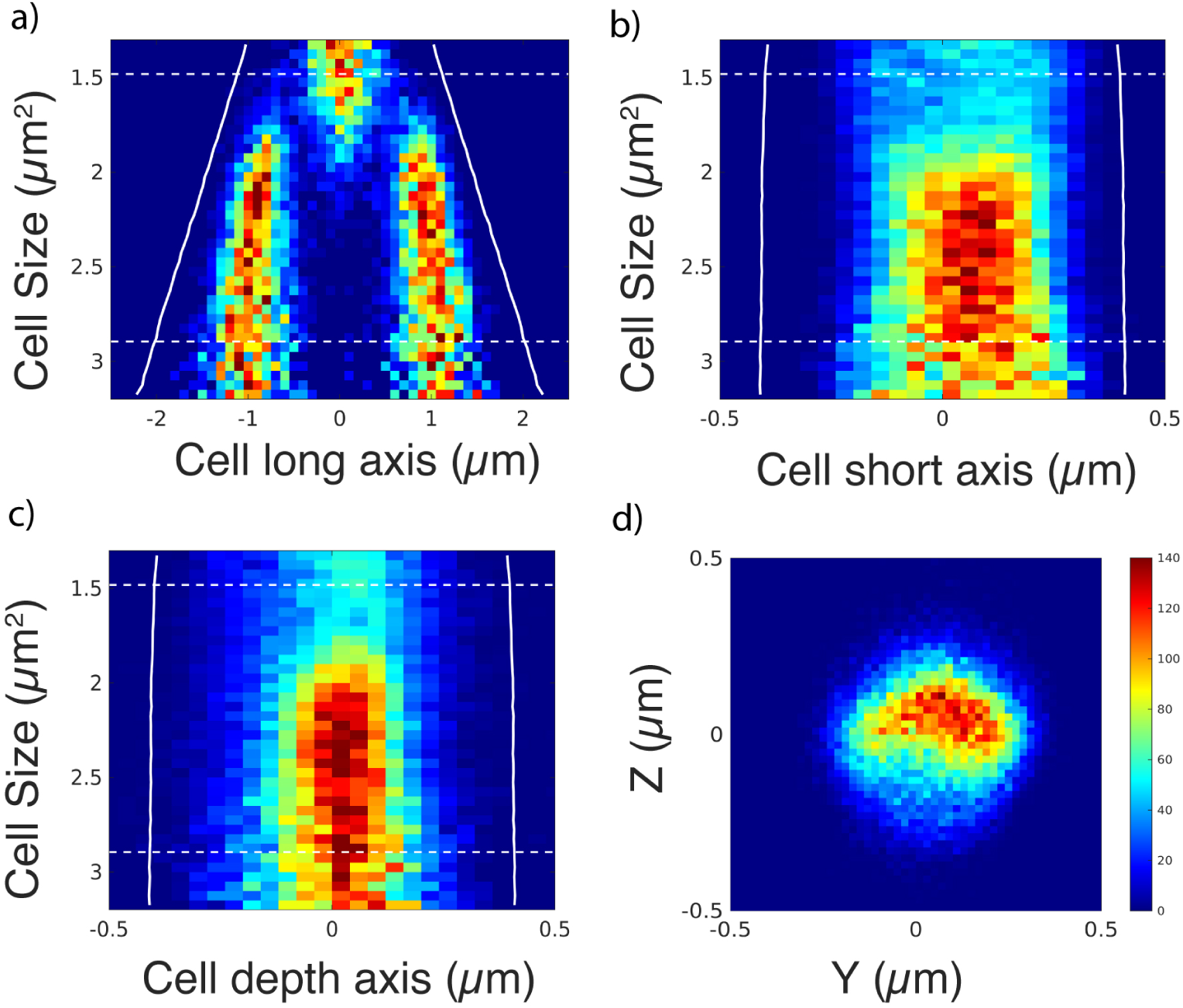
Histograms of emitter locations as a function of cell size for replisome-labeled cells. a) Cell long axis (x-coordinate) vs cell size in *µm*^2^ b) Cell short axis (y-coordinate) vs cell size in *µm*^2^ c) Cell depth axis (z-coordinate) vs cell size in *µm*^2^ d) YZ histogram of all positions. The bin size for the YZ histogram is 20 nm x 20 nm. The color bar represents the number of localizations per bin in this plot.

## Discussion

Live-cell imaging in microbiology has been revolutionized by the use of microfluidics, allowing long experiments with exponentially growing bacteria. Intracellular labeling on these timescales usually calls for endogenously produced fluorescent protein fusions, as any externally added synthetic dye would get diluted by growth. 3D localization of FPs in bacteria has been challenging for a number of reasons, but mainly because of the low signal-to-background ratio of these relatively weak fluorophores. Additional challenges include accurate x, y, and z-localization of the bacterium itself such that the location of the fluorescence emitter can be put in the cell’s internal coordinate system. To map localizations into a cell’s internal coordinate system in XY we use cell segmentation masks. To get the z height accurately in the internal cell coordinates, we use a fluidics chip where the cells are trapped between the PDMS structures and the glass coverslip, at a well-defined distance from the objective. In this paper, we have described how we have overcome the challenges of FP localization in *E. coli*.

To achieve accurate super-resolved 3D localization of emitters, we build on the FD-DeepLoc algorithm. Our most significant addition to the method is obtaining ground-truth training data, which is needed for the algorithm to give meaningful results. Importantly, the cell background has spatial correlations that vary depending on the distance to the cell boundary. Due to the low signal-to-background ratio regime of the experiments and the overlapping spatial dimension of cell background and emitter PSF, removing the structured background is critical for effective use of localization methods. Other higher-order correlations between pixels at different distances to the cell boundary require more complicated modeling and could be captured with more advanced generative modeling techniques.

To test the functionality of the 3D localization method, we tracked distributions of different chromosomal loci and the *E. coli* replisome in 3D over the cell cycle. Since the *E. coli* cell is radially symmetric, we expected the same spatial distribution of emitters along the width and depth of the cell. We could show this radial symmetry for all chromosome and replisome labels. Interestingly, we also observed that some chromosomal loci are located in the periphery of the nucleoid, which has not been observed before.

## Methods

### Bacterial strains

Imaging experiments were performed as described previously [30]. All experiments were performed in M9 minimal medium with 0.06× Pluronic F-108 (Sigma-Aldrich 542342), 0.4 % succinate, and 1× RPMI 1640 amino acid solution (Sigma) at 30 °C. Inoculation of bacterial strains from cryo-stocks into growth medium was performed one day before each experiment. After growing the strains overnight at 30 °C in a shaking incubator (200 rpm), they were diluted 1:100 and loaded in a microfluidic chip. Genotypes of all strains used are described in SI table A1.

### Optical setup

Imaging was performed with a Ti-E (Nikon) microscope equipped with a 100X immersion oil objective (Nikon, NA 1.45, CFI Plan Apochromat Lambda D MRD71970) for both phase-contrast and widefield epi-fluorescence microscopy. Fluorescence images were acquired using a Kinetix sCMOS camera (Teledyne Photometrics). A 515 nm laser (Fandango 150, Cobolt) was used for SYFP2 excitation. Images were acquired at 150 ms exposure time and 5 *W/cm*^2^ power density every 2 min. The excitation laser was triggered by the camera acquisition with a function generator (Tektronix).

Phase-contrast images were acquired with a 50 ms exposure time using a DMK 38UX304 camera (The Imaging Source). The light source used for phase contrast was a 480 nm LED and a TLED+ (Sutter Instruments). The transmitted light was passed through the same FF444/521/608-Di01 (Semrock) triple-band dichroic mirror as the fluorescence and reflected away from the fluorescence light path using a Di02-R514 (Semrock) dichroic mirror. A lens relay system with a phase ring at the back focal plane was used to direct the light onto the camera.

### Microfluidic chip

All microfluidic experiments were done using modified mother-machine devices [9], where cells were loaded into the growth channels and continuously supplied with growth media. Each imaged FOV contained 20 growth channels, corresponding to a 69 *µ*m x 69 *µ*m area.

### Segmentation masks generation

Phase-contrast images were segmented using Omnipose [32, 33] cell segmentation networks trained specifically for mother-machine devices. The segmentation network is described extensively in [32]. These segmentation masks were used to generate EDT to the boundary field whenever required. Values of EDT were rounded to the nearest integer. Cell masks were used for sampling emitter coordinates when simulated microscopy images were provided as input during localization network training. Conversion of global emitter coordinates with respect to the origin of the image into the internal coordinates of each cell was also done based on the cell segmentation masks.

### PSF calibration and CSpline model fitting

Images of beads (TetraSpeck T7279 100 nm size) immobilized on a cover-slip were imaged in 25 nm steps in z and used to generate a PSF using Super-resolution Microscopy Analysis Platform (SMAP)[40]. About 50 beads were localized and aligned in 3D space to generate a mean PSF image stack. All images in the mean PSF stack were also normalized by the maximum intensity value on the mean image of the central three image planes around the focal point of the 3D PSF stack. After normalization, a cubic spline model (Eq.1) was fitted to the normalized PSF stack, giving 64 spline coefficients per voxel. These coefficients were used to interpolate intensity values inside the voxels of a PSF image stack and generate an image for a given emitter coordinate (x, y, z). Intensity variations in z were not corrected at this spline calibration stage, but sampled PSF images in 2D were normalized to sum to 1 before converting to photons during simulations. The 3D PSF spline model follows the piece-wise cubic spline equation:

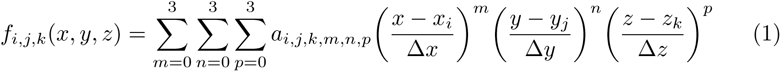

where *f_i,j,k_*(*x, y, z*) gives the normalized 3D PSF intensity around the voxel (i, j, k), Δ*x,* Δ*y* are pixel sizes, and Δ*z* is the step height of the bead stack during acquisition. *x_i_, y_j_, z_k_* are the normalized intensities of the mean bead stack image at the voxel centered at (i, j, k) and *a_i,j,k,m,n,p_* are their cubic spline coefficients. Each voxel has a volume of Δ*x* Δ*y* Δ*z* centered around (i, j, k). The first and second derivatives of the splines of neighboring voxels are constrained to be equal. This function is used to interpolate the PSF for any real-valued location of the emitter coordinate (*x, y, z*).

### Network architecture and training

The neural network models used in the paper consist of two U-nets connected in series [29]. During training, the input to the network consists of fluorescence images simulated using the cubic spline PSF (SI section A.3) and background estimation models (SI section A.5 A.6) and camera coordinate fields defining the sub-region of the camera used at that training step. Emitters were sampled randomly in space with average densities specified per cell. Cell backgrounds for simulations were done using pixel intensity distribution fits described in the SI section A.5 and SI section A.6. Background pixels from the fluidic chip were also sampled from similar fitted distributions. Camera field input was given using the CoordConv strategy [29, 41] to both U-nets to learn camera pixel-dependent noise. A detailed description of the image simulation is described in the SI section A.4.

All networks output 10 channels of the same size as the input image, one channel for probability (*p̂_k_*) of emitter and pixel *k*, 2 channels for sub-pixel offsets (Δ*x̂_k_*, Δ*ŷ_k_*) from the center of each pixel, one channel for axial distance *ẑ_k_* and one for photon counts *I*^^^*_k_*, 4 channels for *σ̂_xk_*, *σ̂_yk_*, *σ̂_zk_*, *σ̂_Ik_* and one channel for camera noise and background free PSFs. Δ*x̂_k_*, Δ*ŷ_k_*, Δ*ẑ_k_* predictions were scaled to be in [−1, 1] using the Tanh activation function while *p̂_k_* and *I*^^^*_k_* were scaled to be in [0, 1]. The maximum photon count was set for each model during training, and the same scaling was applied at the prediction time. All predictions were scaled back to real values (nanometers, intensity values) using appropriate scale factors that were applied on the inputs to the network during training. Photon counts were sampled in the range [750, 3000] and z values were sampled in [−700, 300] nm.

Networks were trained using the AdamW optimizer with a learning-rate scheduler until convergence using PyTorch 2.0. Evaluation metrics used to estimate localization performance were the same as for the SMLM 2016 challenge [34]. Metrics for each training model are shown in the SI section A.8.

### Loss functions

The loss functions used are similar to FD-DeepLoc [29] and consist of four parts.

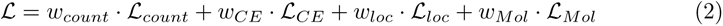

The expected number of emitters in an image with K pixels 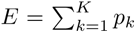, given the Bernoulli detection probability *p_k_* of each pixel, can be approximated as a Gaussian distribution with mean 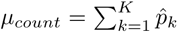 and variance 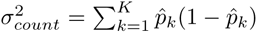, as *p_i_* is independent of *p_j_*for all our experiments.

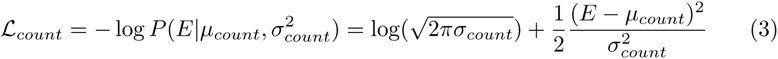

The cross-entropy term *L_CE_* is a simple binary-cross entropy between the ground-truth pixel-wise probability map *p_k_* and the predicted probability map *p̂_k_*:

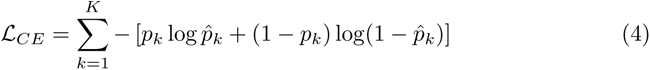

Localization error *_loc_*between true emitter locations and predicted emitter locations is approximated using a Gaussian-mixture model over the predicted per-pixel distributions:

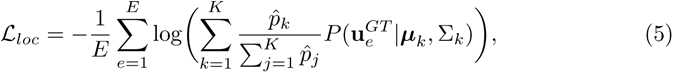

where 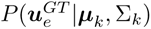 is the probability density of ground-truth values **u** = [*x, y, z, I*] of emitter *e* calculated using the pixel-wise prediction distribution. This distribution is approximated as a 4-d Gaussian distribution with diagonal covariance matrix 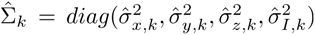 and mean 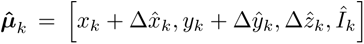 and calculated as

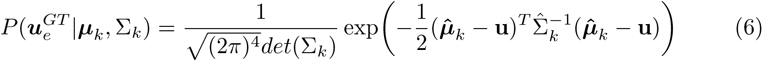

Finally, *L_Mol_*minimizes the error between simulated PSF and predicted PSF images:

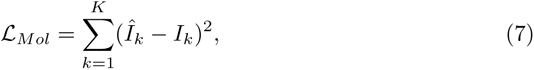

where *I* is the image of noise-free simulated emitters (SI figure A3g). Additionally, each loss component is multiplied with a weight such that all four loss components have the same order of magnitude. The weights of different loss components (*w_count_, w_CE_, w_loc_, w_Mol_*) are set to 0.2, 1.0, 8000, and 0.25 respectively.

### sCMOS camera model

An sCMOS camera (Photometrics Kinetix) was calibrated using the Accent plugin [42] in Fiji to estimate pixel-dependent camera parameters (SI A.1) such as e-/ADU, read noise, and thermal noise. The camera was operated in sensitivity mode (12-bit) and dark images for calibration were acquired with varying exposure times using a Micro-manager 1.4 script. The same sCMOS sub-region was used during camera calibration and when acquiring bead and cell data. Pixel-dependent camera parameters, gain, read noise, and thermal noise, were used to sample noise during the simulation of training data for all neural networks. Due to the noisy estimation of gain values per pixel using this method, the median gain value (~2.58 ADU/e-) was used for conversions between e- and ADU wherever required. Camera parameter distributions are shown in SI methods A.2.

### Inference and Post-processing

Network outputs of experimental images were processed to construct a list of emitters in each image. The probability output channel was used to obtain emitter pixel locations after non-max-suppression as described in [29]. Offsets from the center of the pixel and localization precision were obtained by masking the other output channels using the emitter pixel locations. Final emitter locations were converted to a list of coordinates in nanometers from the origin of the image (top left corner).

Before converting the global coordinates of emitters into internal coordinates inside cells, they were overlaid on cell segmentation masks and filtered for any emitters that fell outside the cell boundary. Cell tracking was performed using the Baxter algorithm [43] and only emitters that fall in tracked cells were used for converting to internal coordinates. XY coordinates of emitters belonging to a cell were transformed into coordinates relative to the cell’s long-axis and short-axes. The z coordinate, however, is not transformed due to a lack of origin reference along the z-axis of the cells. The corrections for variations in the height of the imaging plane were done as described in the main text. For illustration purposes, the boundaries of cells in the depth axis are the same as that of the short axis in Fig 3 and Fig 4.

### Statistics and reproducibility

The results shown for different chromosomal loci and replisome complexes were reproducible across 16 different FOVs.

## Acknowledgments

We wish to thank Irmeli Barkefors for helpful input on the manuscript. We acknowledge funding from SSF (ARC19-0016), the European Research Council (BIGGER:885360), the Knut and Alice Wallenberg Foundation (2016.0077; 2017.0291; 2019.0439) and the eSSENCE e-science initiative to J.E.

## Author Contributions

J.E conceived the project. P.K developed the method and put together the processing pipeline. K.G performed all the experiments. S.Z wrote the code for image transformations and 3D localization corrections. E.A built the optical setup and provided feedback during the entire project. P.K, K.G, J.E wrote the paper with inputs from all co-authors.

## Data and Code availability

Code for 3D localization will be available at https://www.github.com/karempudi/cellbgnet.git upon publication. Raw datasets, neural network models, and code for 3D distribution plots will also be made publicly available upon publication.

## Appendix A Supplementary Information

### A.1 sCMOS camera model

Camera noise model is approximated using experimentally determined parameters (SI A.2) and is used to convert photons to image units (ADU) and vice-versa. Photon statistics, pure light intensity fluctuations, is modeled as a Poisson process. Conversion from photons to electrons by absorption of photons in the sCMOS sensor is accounted for by an absorption probability that scales with the mean of the Poisson process.

Different sources of noise considered are shot noise, read-noise (RN) and thermal-noise (TN). Shot noise accounts for Poissonian distribution of photons hitting the camera pixels. Read-noise accounts for noise in reading electrons, and thermal noise accounts for electrons generated due to thermal excitation.

If *λ_ph,k_*are the expected number of photons collected in pixel *k*, and *qe* is quantum efficiency at a given imaging wavelength, then the expected number of electrons from pixel k, is

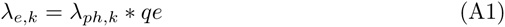

During any given camera exposure the amount of electrons *s_k_* accumulated in a pixel can be modeled according to equation (A2). The signal observed at pixel *k*, *s_k_* follows a Poissonian distribution with mean given by *λ_e,k_*,

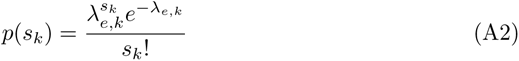

Noise in reading electrons and from thermal electrons for sCMOS cameras are typically modelled as Gaussian distribution with variance 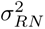 and 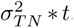, where *t* is exposure time in seconds. In our modeling we estimate them using dark-noise images (SI A.2) and combine them into one Gaussian distribution with variance 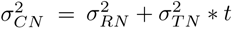. Camera noise electrons are sampled from the Gaussian,

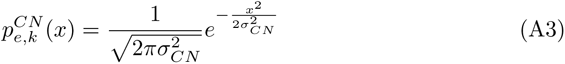

and are added to *s_k_*. Gain (ADU/e-) is used to convert electron signal to counts (ADU) and a predefined offset value is added to the counts to obtain final image (*ADU_k_*).

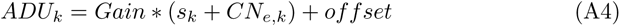

### A.2 Camera calibration and camera noise sampling

5000 dark images at varying exposure times of 25 ms, 50 ms, 75 ms, 150 ms, 300 ms over the region-of-interest (ROI)(1041 x 1302 pixels) on the camera (3200 x 3200 pixels) were used to calibrate baseline, gain, pixel-wise read noise and thermal noise parameters using the Accent plugin [42] and Fiji. During simulations of training data, read-noise (RN) and thermal noise (TN) were combined into one noise-map defining pixel-wise standard deviation for the particular exposure time used of imaging experimental data. Camera noise is sampled for each pixel using this standard deviation map. Fig A1 show the distributions of gain and combined read-noise and thermal noise. Due to the noisy nature of gain-estimation using this method, we use median gain (*∼* 2.58) of all conversions between e- and counts (Fig A1).

**Fig. A1.**
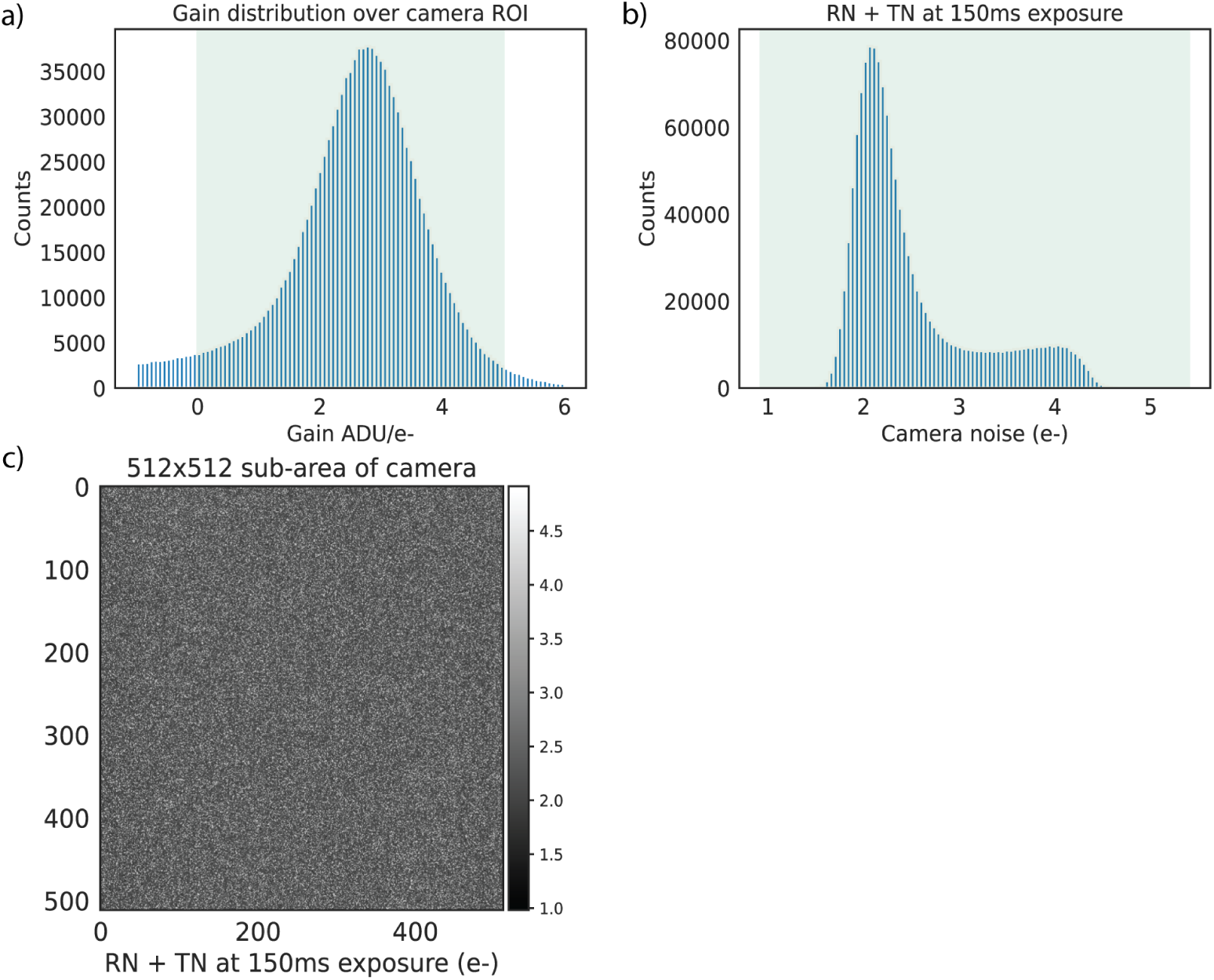
Camera gain, camera noise histograms and maps of camera noise. a) Gain distribution over the ROI b) Read noise+thermal noise standard deviation distribution at 150 ms exposure. c) Read noise + thermal noise over a smaller sub-region on the full ROI in e-.

### A.3 Experimental astigmatic point spread function sampling

The mean PSF for each fluorescence channel was obtained from aggregating ~ 50 beads stacks and aligning them using 3D cross-correlation [40]. The stack was normalized using the peak intensity in the mean image of 3 frames around the focal point of the PSF (z = 0). After fitting a spline model to the normalized mean PSF stack, images of the PSF were sampled at the center pixel of a 41x41 image and normalized such that the image sums to 1 for each z sampled. Images of the PSF are shown in Fig A2. Fitting of splines was done using SMAP [40] in MATLAB to obtain coefficients which were later used for sampling emitter images using DECODE’s spline sampling function in Python. Fig A2 b also shows the Cramer-Rao lower bound (CRLB) of the localization precision achievable using the model of the PSF in the low noise regime with a uniform background model (1875 photons for the emitters and 20 photons for the background). When we do experiments we confine ourselves to operating in the range -700 nm to 300 nm.

**Fig. A2.**
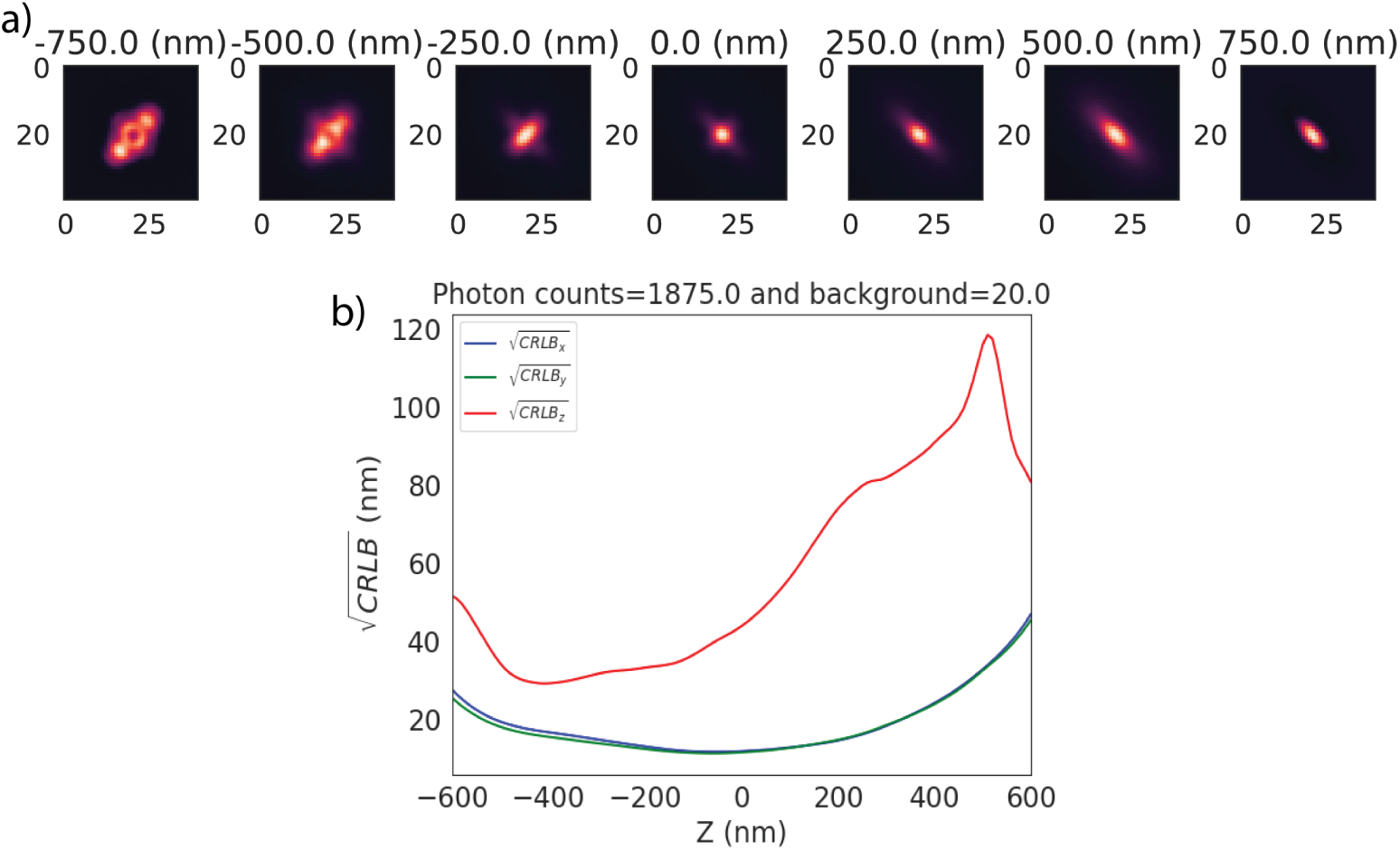
a) Normalized PSF function stack in the YFP imaging channel obtained from averaging 50 beads. Each image of the PSF is sampled from cspline model where emitters have varying z from -750 nm to +750 nm placed at the center of the image and normalized such that each image sums to 1.0. b) CRLB of the localization precision in x, y and z using the spline model.

### A.4 Emitter sampling using cell segmentation and background distributions

Phase-contrast images (1041 x 1302 pixels) are used to obtain segmentation masks using the Omnipose method previously described in [32]. Average cell area is calculated over 300 images and is used to estimate probability of an emitter in a cell pixel, assuming an average of 2 emitters per cell. Random crops of size 128 x 128 are sampled from the segmentation masks. Centers of pixels containing emitters are sampled using binomial distributions of the probability map. XY offsets of the emitters from center of the pixel are uniformly sampled between [-0.5, 0.5], while z is sampled uniformly between [-700, 300] nanometers. PSF images of size 41x41 pixels are sampled using a spline model (SI section A2) and normalized to 1 before multiplying with photon counts. Photon counts are uniformly sampled from defined range to match ADU levels of experimental data. Each emitter image is then placed on a larger image 128x128 pixels.

Cell segmentation masks are converted to EDT and background photons are sampled using gamma distributions specific to the experimental data (SI section A.5, A.6). Emitters’ PSF image is added to sampled background photons images and passed through the camera noise model to generate final simulated image. Camera noise is sampled from a Gaussian distribution whose sigma is calculated per pixel during camera calibration. As we only use a smaller region (128x128 pixels) of the larger area of the camera chip, the locations of the smaller regions are encoded using the Coord-Conv strategy [29, 41] and are used as inputs to the network. Fig A3 shows a series of images that illustrate the image generation process.

**Fig. A3.**
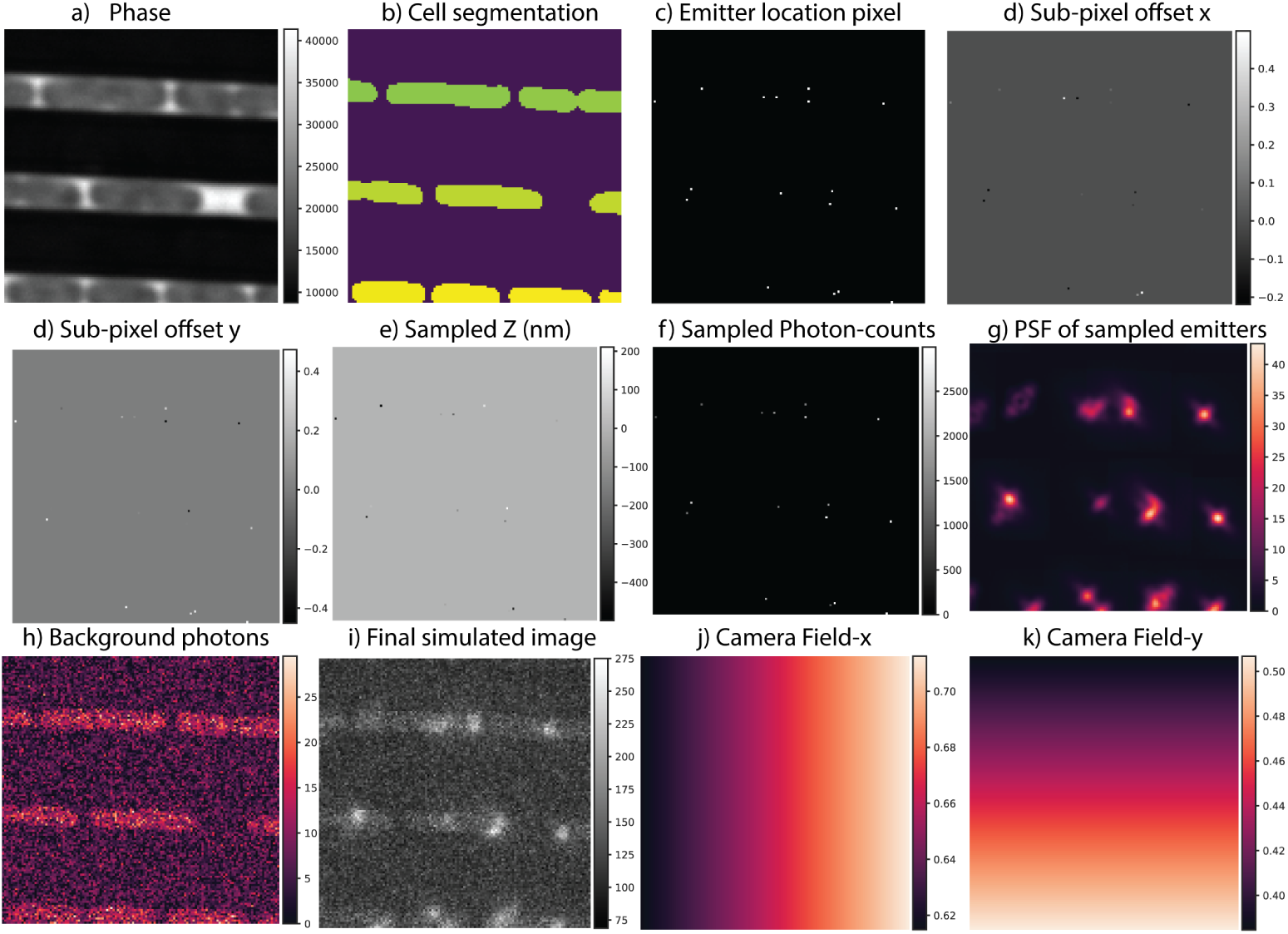
Emitter sampling and simulation of image generation of 128x128 image a) Phase-contrast image b) Cell segmentation mask c) Sampled emitter pixels’ centers d-g) Sampled x, y, sub-pixel offsets from the center of the pixel. z position sampled in range [-700, 300]nm, photon counts in the range [750–3000] g) Photons of all emitters overlayed onto a single image h) Sampled background photons from gamma distributions of background photons i) Final images with emitter and background photons added and converted to ADU after adding pixel-dependant camera noise j-k) XY field of the sub-region on the camera ROI with origin at [800, 400]. Camera ROI is of size 1302x1041 pixels

### A.5 Cell background estimation for chromosome dots

Phase-contrast images are segmented to obtain cell segmentation masks after transforming phase images to the same size as fluorescence images. Cell segmentation labels are dilated by 1 pixel. Pixels on each image are masked using cell segmentation into two categories, inside and outside the cells. Pixels outside cells are used to estimate gamma distribution of background photons coming from the chip (SI Fig A5a). Pixels inside the cells are used to estimate background distributions as a function of distance to the boundary of a cell. Inside cells, only pixels less than the 75th percentile are used to estimate background, removing most potential emitters (SI Fig A4). Collections of pixels from each images at varying distances from the boundary are used to fit gamma distributions, whose mean and variance are used to sample realistic cell backgrounds during simulations. Fig A5b shows fits of data and distributions. This procedure is repeated over 300 images and variation of means of the fitted distributions is shown in Fig A5c. From this, we conclude that background can be approximated as a function of distance from the boundary. Mean and variance masks are created for each cell mask used to sample emitters using the average values over all 300 images. Fig A5d shows these maps for an example cell mask. Sampled background ADU is converted to photons and is used for simulations (Fig A3h).

**Fig. A4.**
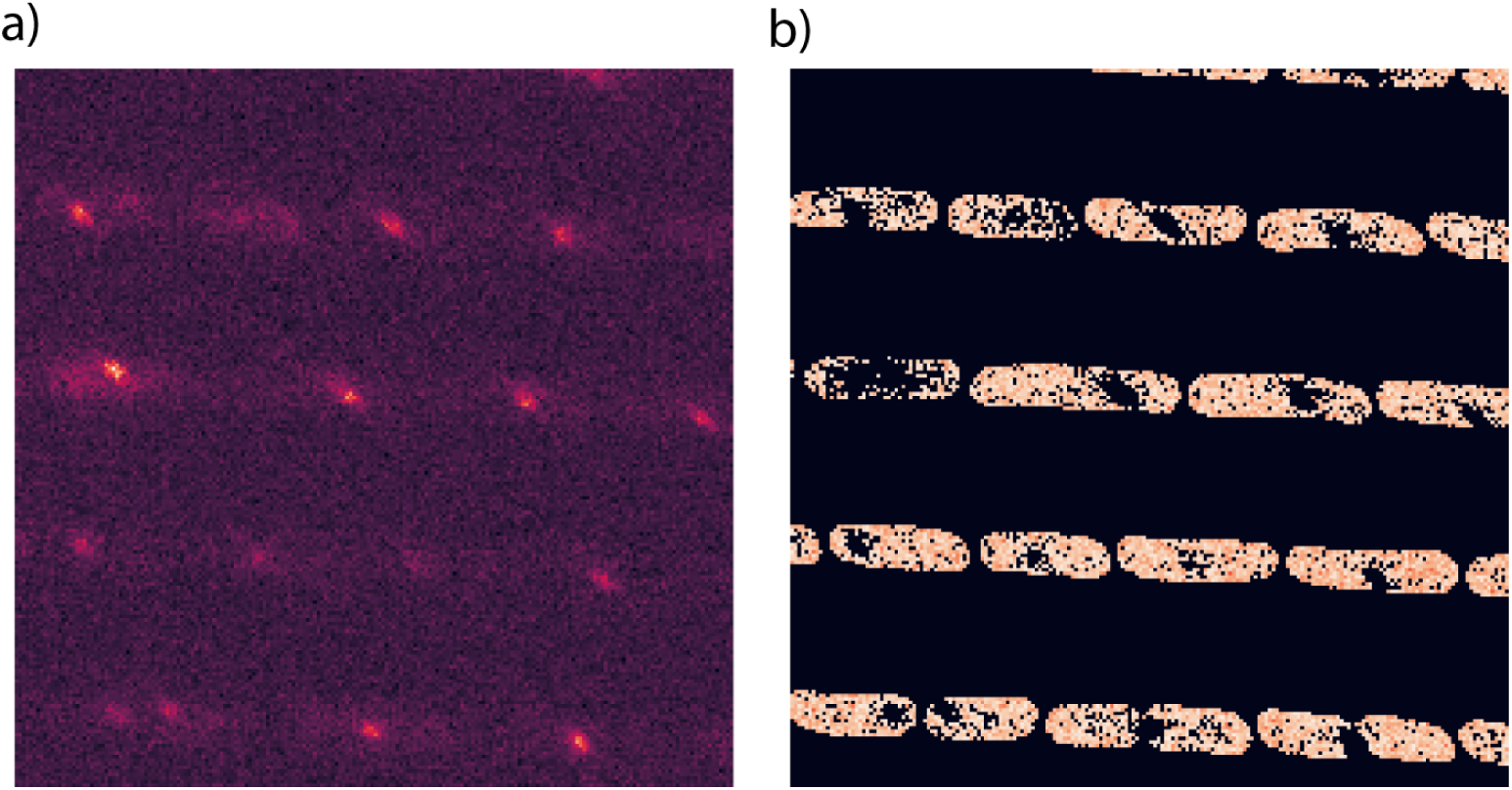
a)Fluorescent image with emitters b) Image with emitters removed to estimate cell background

**Fig. A5.**
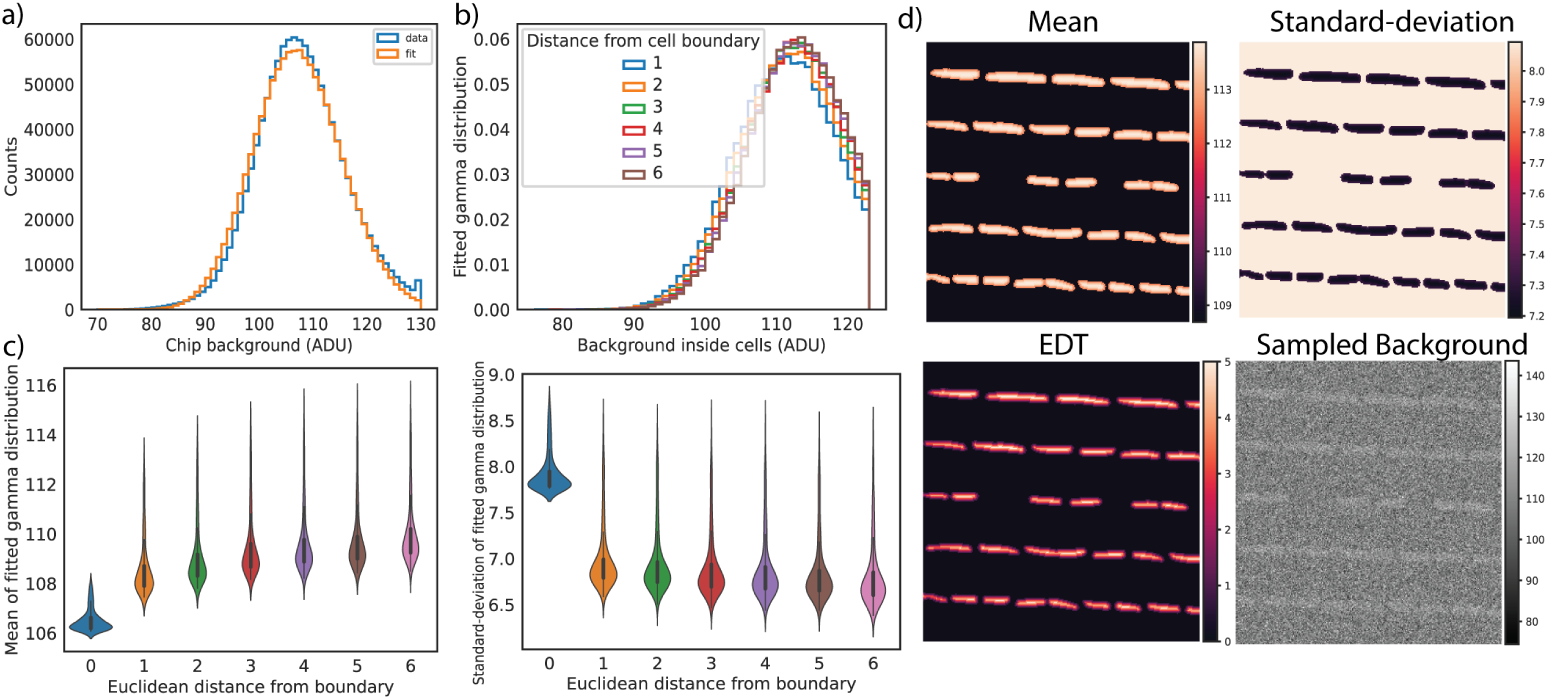
Cell and fluidic chip background for chromosome dots a) Chip background distribution and fit over one image b) Distribution fits as a function of Euclidean-distance form the boundary (EDT) for pixels inside cells c) Mean and standard deviation of the fitted gamma distribution over 300 images as a function of EDT. EDT=0 corresponds to chip background. d) Mean and variance over an example cells mask and corresponding sampled background using distributions from c.

In the astigmatic PSF (Fig A2) simulated with emitters at the center of the frame, the brightest pixel at 0 nm has ~ 2.1 % of the photons while -500 nm and +500 nm has ~ 0.5 %. Using this information together with ADU values observed in experimental images, photon count range for the simulations are set between 750 and 3000 photons.

### A.6 Cell background estimation for replisome dots

A procedure similar to the one described in the previous section was repeated for the replisome labelled cells. The key difference between the data sets is the higher background noise in this data set (SI Fig A6c) compared to the chromosomal labels (SI Fig A5c)

**Fig. A6.**
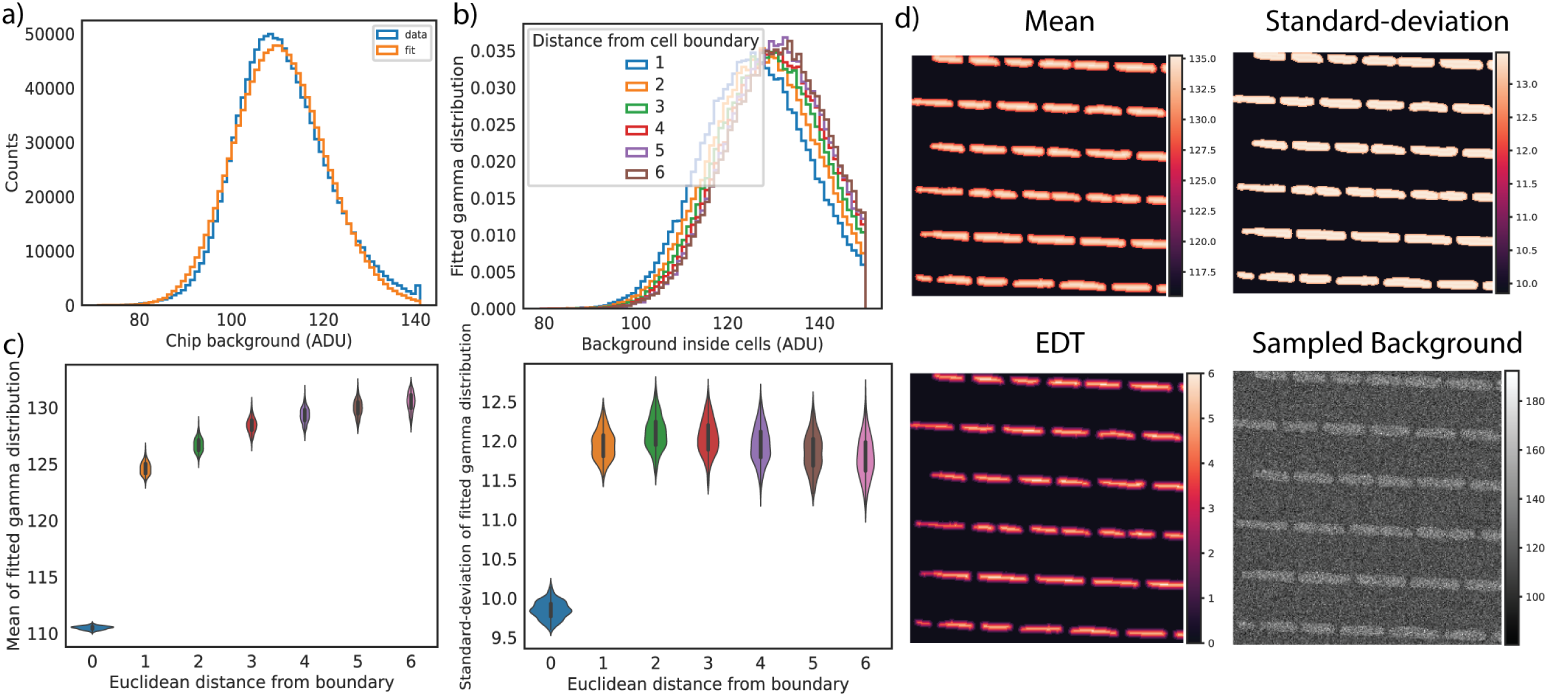
Cell and chip background for replisome dots a) Chip background distribution and fit over one image b) Distribution fits as a function of EDT for pixels inside cells c) Mean and standard deviation of the fitted gamma distribution over 300 images as a function of EDT. EDT=0 corresponds to chip background. d) Mean and variance over an example cells mask and corresponding sampled background using distributions from c.

### A.7 Training hyper-parameters

The localization networks were trained for 20000-40000 iterations with an initial learning rate (0.0006) that was reduced by 90% after every 5000 iterations. The batch size was 64. AdamW optimizer with a weight decay of 0.1 was used for training.

### A.8 Network training performances and evaluations of trained models

For an input image of size 128x128 pixels containing emitters and corresponding camera XY-field, the dual U-net predicts probabilities, XY sub-pixel offsets, z and photon counts along with their *σ*’s and PSF of all predicted emitters in the image. These values are scaled to appropriate ranges in which the model was trained i.e nanometers for x, y, z and real values for photon counts. Model predictions are matched to ground-truth emitters after thresholding network outputs and doing non-max-suppression on probabilities. Each emitter is matched to ground truth emitters in radius of 250 nm around the predicted localization. A total of ~ 21000 emitters simulated in 30 evaluation images of full camera FOV size 1041x1302 pixels were used to calculate these metrics during training of network. A final precision ~ 1.0 and recall of ~ 0.8-0.95 was achieved after 20,000-40,000 iterations of training with a batch size of 64 per iteration for both chromosomal loci dots and replisome dots.

Fig A7 shows the convergence of the loss function, RMSE-x, RMSE-y, RMSE-z, precision (TP / TP + FP) and recall values (TP / TP + FN) for one model trained for inferring localizations of chromosomal loci labelled data.

**Fig. A7.**
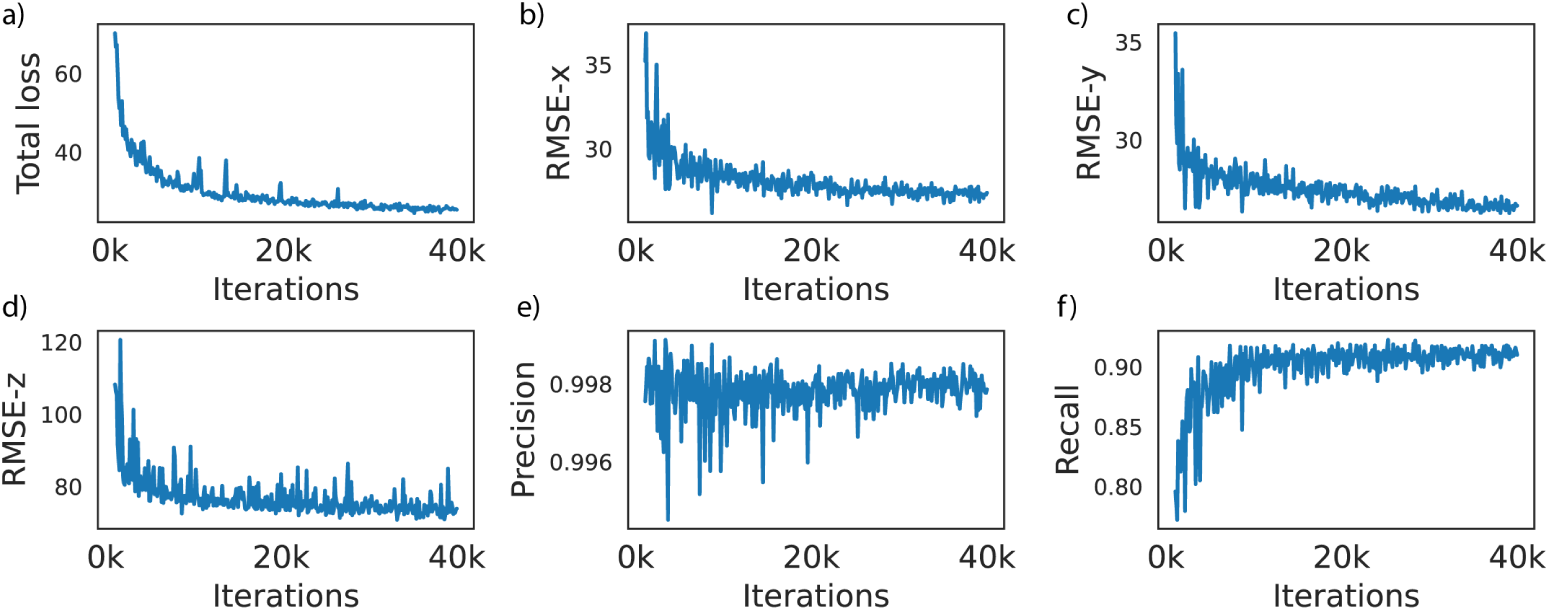
Network performance metrics of a model trained in range [-500, 500] nm of the PSF as function of training-loop iteration number a) Overall loss value b) RMSE-x c) RMSE-y d) RMSE-z e) Precision (TP / TP + FP) f) Recall (TP / TP + FN).

Evaluation of any biases in learning requires looking a distributions of residuals of the simulated ground-truth emitters and model predictions. Fig A8a-d shows four distributions for deviations of x, y, z and photon count predictions from the matched ground-truth emitters. Sub-pixel offset prediction distributions are shown in Fig A8e-f. Means of deviations of localizations predicted by the network are close to 0, indicating that the predictions are not biased in the overall z-range over which the network is trained on.

**Fig. A8.**
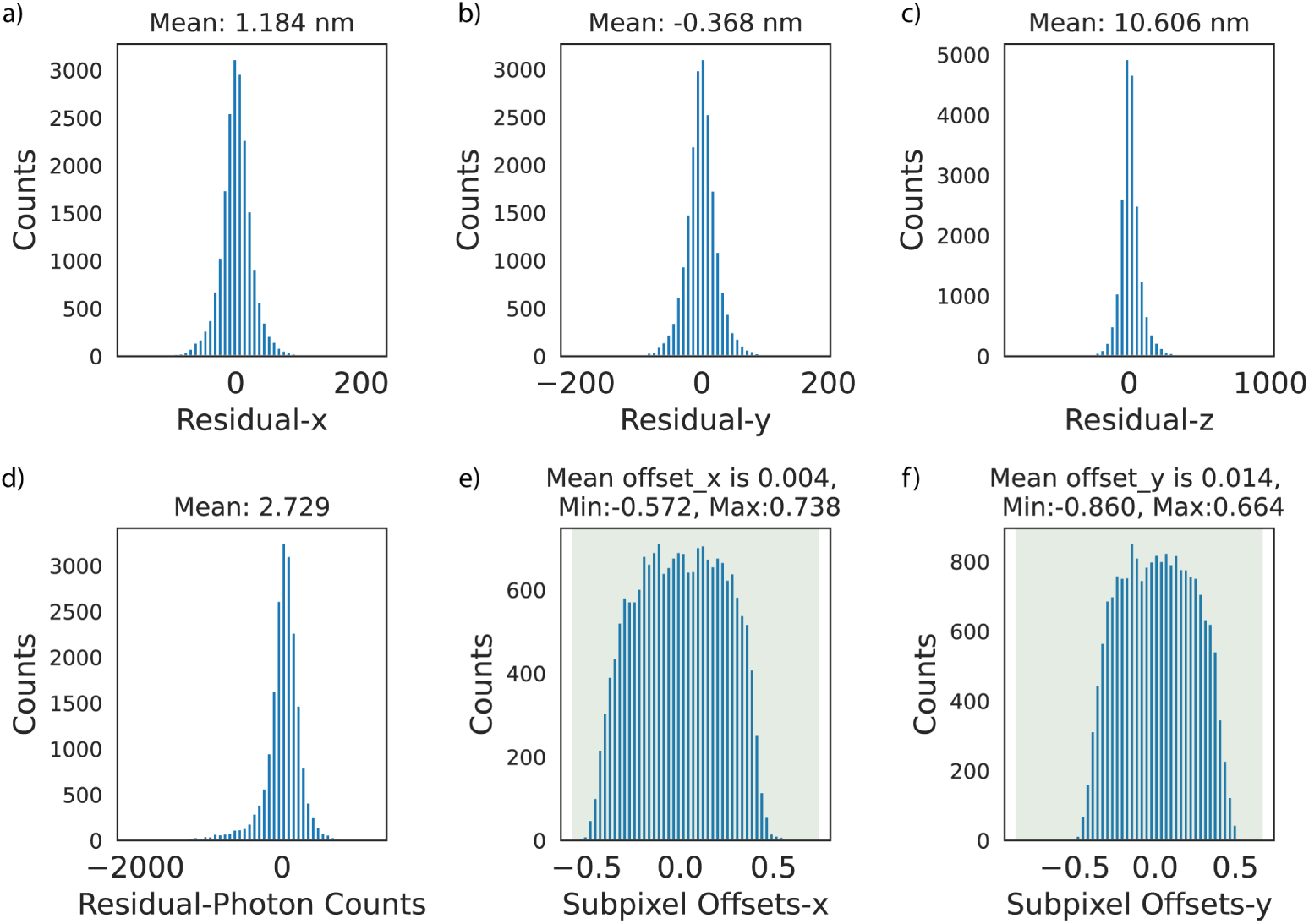
Distributions of deviations between matched ground-truth emitter locations and localizations predicted by the model. a) x-deviation b) y-deviation c) z-deviation d) photon count deviation e) Sub-pixel offset-x distribution f) Sub-pixel offset-y distribution

### A.9 Transformations between phase and fluorescence

Images of fluorescent beads (TetraSpeck T7281 500 nm size) were captured on both cameras. Bead positions were selected manually on both images using the *cpselect* function in MATLAB. Geometric transformations of the projective kind were fit using the *fitgeotrans* function in MATLAB, similar to the procedures described in [30].

### A.10 Additional distributions of chromosomal loci data

The localizations used in Fig 3 where chromosomal loci labelled at different positions also have associated uncertainties (*σ_x_*, *σ_y_* and *σ_z_*) predicted by the network. These uncertainties are plotted against the internal coordinate assigned to each localization in x, y and z. The internal coordinate in x is scaled in the range [0, 1], while that of y is in [-1, 1] to remove the effects of variation in cell-size. We expect a symmetric distribution around 0.5 for internal-x and 0 for internal-y. For the z distribution however, we expect it to be symmetric around 0 nm, if the localization precision is uniform across the operating range in height. SI Fig A9 show these distributions and they follow expected patterns.

**Fig. A9.**
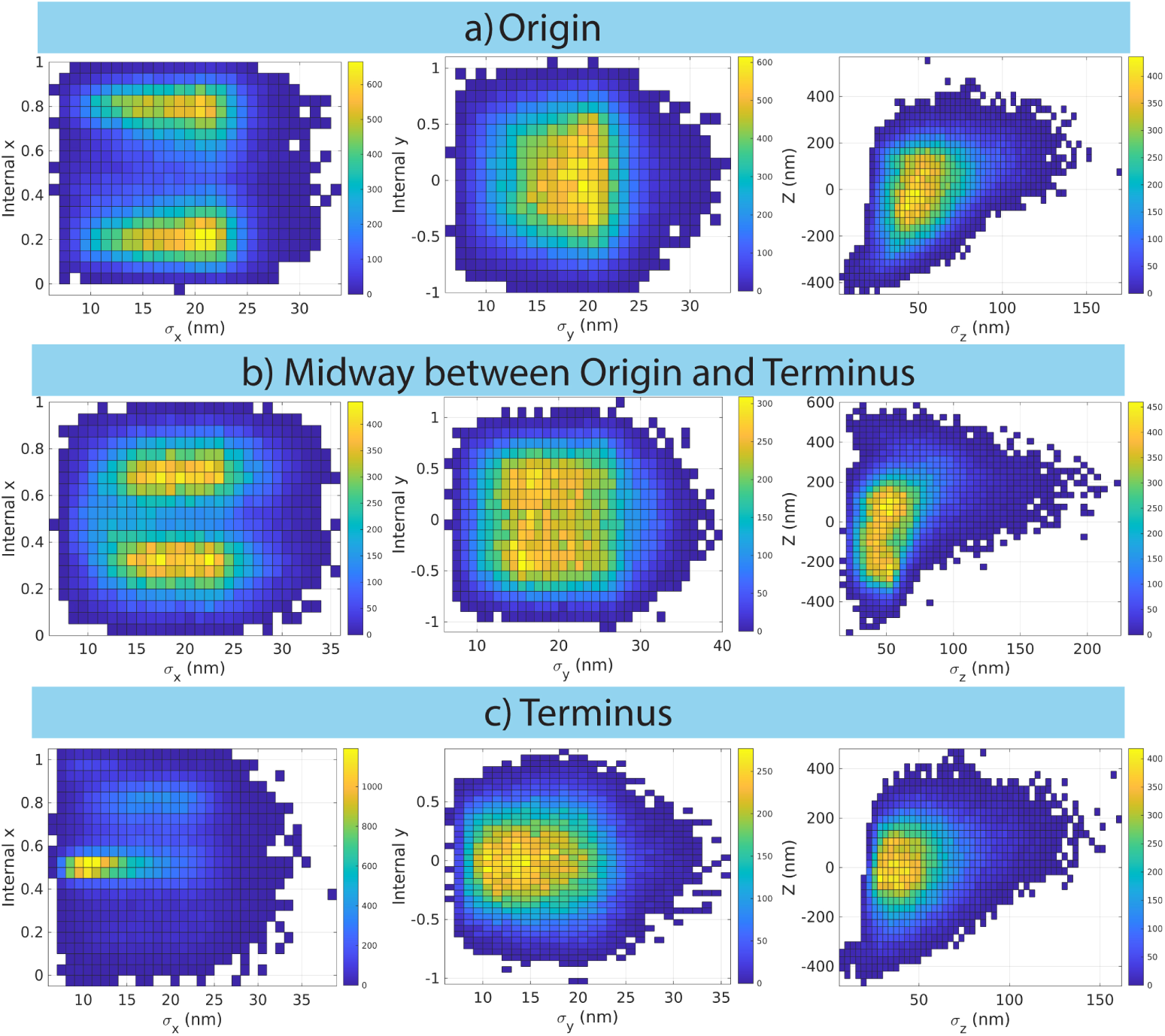
Uncertainties of emitter localizations as functions of cells’ internal coordinates for x and y and real z coordinate for different chromosomal loci. The internal x coordinate is scaled to the range [0, 1], while the internal y is in the range [−1, 1]. a) Origin b) Midway between origin and terminus c) Terminus.

### A.11 Additional distributions of replisome data

Uncertainties of localizations for the replisome data plotted in Fig 5 are shown in SI Fig A10. They follow a similar expected pattern described in the previous section.

**Fig. A10.**
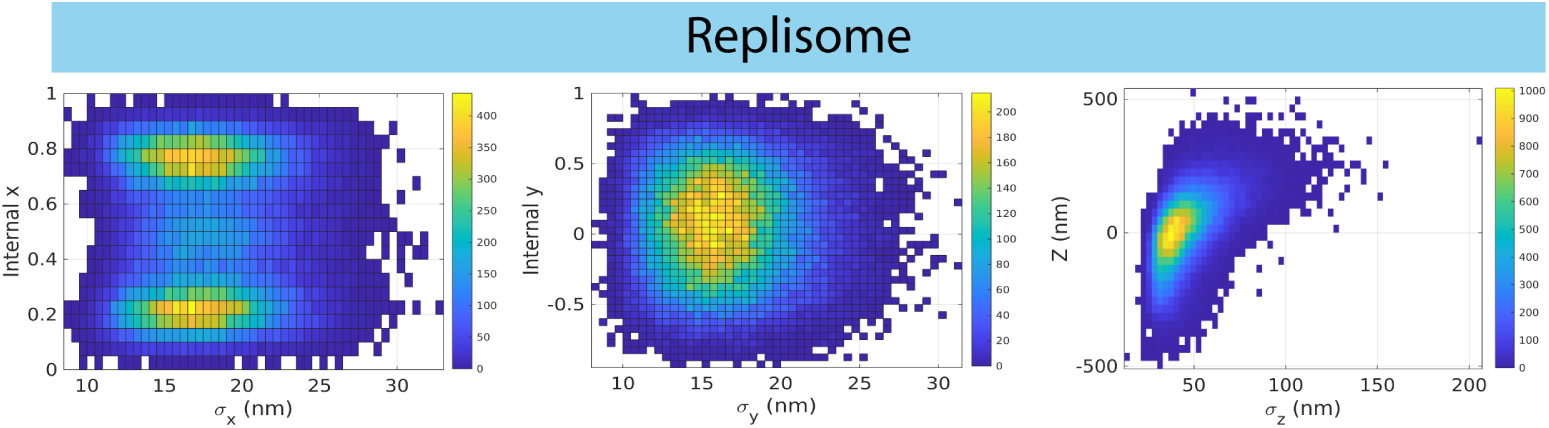
Uncertainties of emitter localizations as function of cells’ internal coordinate for x and y and real z coordinate for replisome complex label. The internal x coordinate is scaled to the range [0, 1], while the internal y is in the range [−1, 1].)

### A.12 Bacterial strains

**Table A1.**
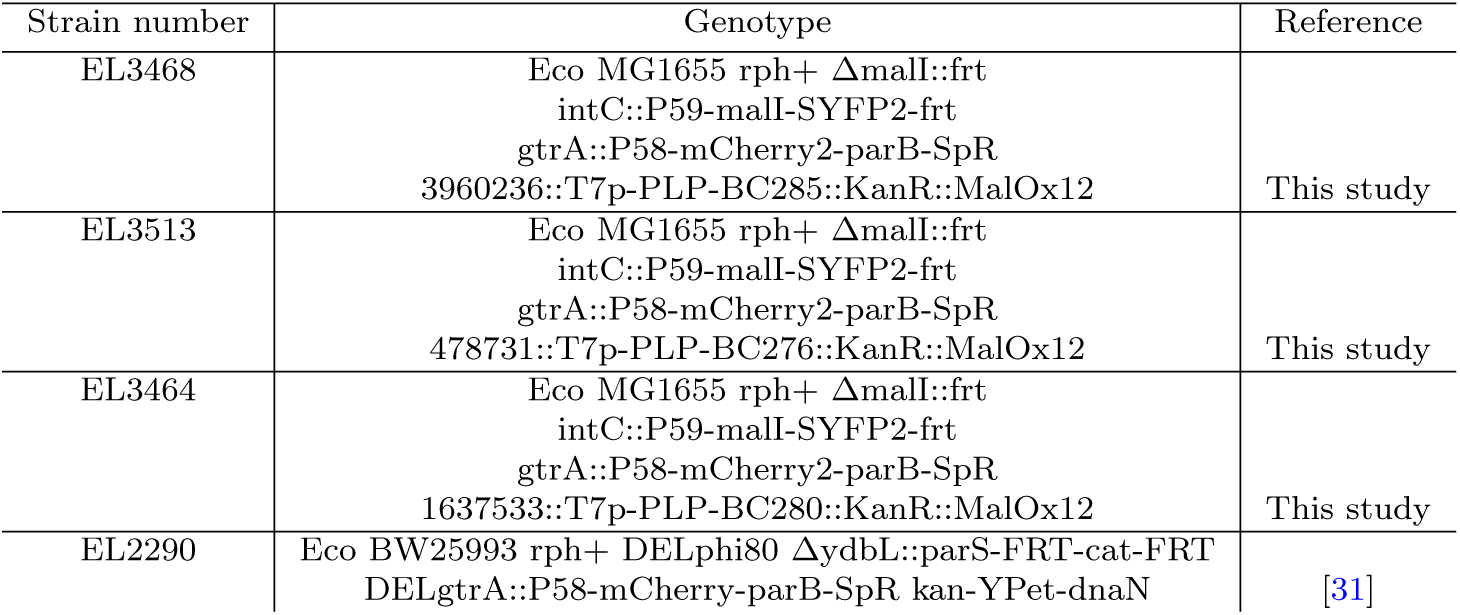
List of bacterial strains used in this study. For the chromosomal labels the number indicates the chromosomal base pair position where the label has been introduced.

